# Formal axioms in biomedical ontologies improve analysis and interpretation of associated data

**DOI:** 10.1101/536649

**Authors:** Fatima Zohra Smaili, Xin Gao, Robert Hoehndorf

## Abstract

**Motivation:** There are now over 500 ontologies in the life sciences. Over the past years, significant resources have been invested into formalizing these biomedical ontologies. Formal axioms in ontologies have been developed and used to detect and ensure ontology consistency, find unsatisfiable classes, improve interoperability, guide ontology extension through the application of axiom-based design patterns, and encode domain background knowledge. At the same time, ontologies have extended their amount of human-readable information such as labels and definitions as well as other meta-data. As a consequence, biomedical ontologies now form large formalized domain knowledge bases and have a potential to improve ontology-based data analysis by providing background knowledge and relations between biological entities that are not otherwise connected.

**Results:** We evaluate the contribution of formal axioms and ontology meta-data to the ontology-based prediction of protein-protein interactions and gene–disease associations. We find that the formal axioms that have been created for the Gene Ontology and several other ontologies significantly improve ontology-based prediction models through provision of domain-specific background knowledge. Furthermore, we find that the labels, synonyms and definitions in ontologies can also provide background knowledge that may be exploited for prediction. The axioms and meta-data of different ontologies contribute in varying degrees to improving data analysis. Our results have major implications on the further development of formal knowledge bases and ontologies in the life sciences, in particular as machine learning methods are more frequently being applied. Our findings clearly motivate the need for further development, and the systematic, application-driven evaluation and improvement, of formal axioms in ontologies.

**Availability:** https://github.com/bio-ontology-research-group/tsoe

## 1 Introduction

Biomedical ontologies are widely used to formally represent the classes and relations within a domain and to provide a structured, controlled vocabulary for the annotations of biological entities (Smith *et al.*, 2007). Over the past years, significant efforts have been made to enrich ontologies by incorporating formalized background knowledge as well as meta-data that improve accessibility and utility of the ontologies (Smith *et al.*, 2007; Mungall *et al.*, 2011). Incorporation of formal axioms contributes to detecting whether ontologies are consistent (Smith *et al.*, 2003; Smith and Brochhausen, 2010; Stevens *et al.*, 2003), enables automated reasoning and expressive queries (Hoehndorf *et al.*, 2015a; da Silva *et al.*, 2017; Jupp *et al.*, 2012), facilitates connecting and integrating ontologies of different domains through the application of ontology design patterns (Osumi-Sutherland *et al.*, 2017; Hoehndorf *et al.*, 2010), and can be used to guide ontology development (Köhler *et al.*, 2013; Alghamdi *et al.*, 2018). While axioms are mainly exploited through automated tools and methods, ontologies also contain labels, synonyms, and definitions (Hoehndorf *et al.*, 2015b); improving the human-accessible components of ontologies has also been a major focus of ontology development (Köhler *et al.*, 2006); for example, including “good” natural language definitions and adequate labels is a requirement for biomedical ontologies in the Open Biomedical Ontologies (OBO) Foundry (Smith *et al.*, 2007), an initiative to collaboratively develop a set of reference ontologies in the biomedical domains.

The amount of information contained in ontologies, and the rigor with which this information has been created, verified, and represented, may also improve domain-specific data analysis through the provision of background knowledge (Garcez and Lamb, 2004). Domain-specific background knowledge can limit the solution space in optimization and search problems (Garcez and Lamb, 2004; Besold *et al.*, 2017; Garcez *et al.*, 2015) and therefore allow finding solutions faster.

The Gene Ontology (GO) (Ashburner *et al.*, 2000) is a biomedical ontology that formally represents several aspects of biological systems, in particular the molecular functions that gene products may have, the biological processes they may be involved in, and the cellular components in which they are located (Huntley *et al.*, 2014b). The GO has been extensively used to provide annotations to gene products through a combination of manual curation of literature and electronic assignments created using algorithms based on sequence similarity, keywords, domain information, and others (Huntley *et al.*, 2014a). Databases such as the GO Annotation (GOA) database (Huntley *et al.*, 2015) use GO to annotate more than 50 million proteins (Huntley *et al.*, 2015).

Due to its central role and importance in molecular biology, significant resources have been invested in the development of GO. Over the years, substantial efforts have been made to improve the coverage of GO through the addition of new classes (Consortium, 2014, 2016). In addition to new classes, GO has also been extended through axioms that characterize the intended meaning of a class formally (Mungall *et al.*, 2011). Specifically, GO now includes links between GO classes and classes in other biomedical ontologies (Bada and Hunter, 2008) in an extended version of GO (which we refer to as “GO-Plus”) (Consortium, 2014, 2016). These axioms are particularly useful in keeping GO complete and logically consistent with other ontologies as well as in guiding ontology development (Consortium, 2016; Bodenreider and Burgun, 2005; Johnson *et al.*, 2006; Mungall *et al.*, 2011). There are now more than 90,000 inter-ontology axioms in GO-Plus that weave GO together with several other ontologies in the biomedical domain.

While these axioms have primarily been developed to tackle the problem of continuously developing GO while maintaining consistency (within GO and other ontologies) as well as to maintain biological accuracy, they also have the potential to significantly improve GO-based data analysis by introducing new associations between classes that are not present in GO but arise through information in other, related ontologies. For example, the GO class *Histidine catabolic process to glutamate and formamide* (`GO:0019556`) and the GO class *Formamide metabolic process* (`GO:0043606`) are not directly (or closely) related in the GO hierarchy but both are related to the ChEBI class *Formamide* (`CHEBI:16397`) through axioms formulated in the Web Ontology Language (OWL) (Grau *et al.*, 2008), a formal language based on Description Logics (Horrocks *et al.*, 2006). If a data analysis method can utilize the axioms in this formal language, we expect improved performance results when applied to different domains.

A task or method that explicitly relies on the axioms or the meta-data in ontologies can not only be used to improve data analysis but also to evaluate the “quality” of axioms in ontologies in contributing to such an analysis task (Hoehndorf *et al.*, 2012). Specifically, such a method would enable determining whether axioms and formalized knowledge contribute to biomedical data analysis, and allow evaluating and comparing how much they contribute to particular tasks.

Recently, several machine learning methods became available that make it possible to utilize different components of ontologies – axioms, labels, definitions, and other kinds of meta-data – in machine learning tasks without the need for manual extraction of features (which may introduce a bias). Here, we use two recently developed techniques, Onto2Vec (Smaili *et al.*, 2018a) and OPA2Vec (Smaili *et al.*, 2018b), to predict protein interactions based on functional information and gene–disease associations based on phenotypes. We evaluate the effect of the axioms that have been added to the GO as well as the effect of adding the axioms of additional domain ontologies as the background knowledge. We demonstrate that the formal axioms that have been created for GO and other ontologies improve predictive data analysis by providing background knowledge about biological domains. Our approach is also applicable to evaluation of meta-data such as labels and definitions and their contribution to predictive analysis of biomedical data. We find that labels and definitions in ontologies can fill gaps in domain knowledge that are not covered by the axioms and further improve prediction; however, the labels and definitions also have the potential to add noise or bias to prediction results. Finally, we also improve the performance of predicting protein interactions and gene–disease associations through ontologies.

Overall, our results demonstrate the value that ontologies provide to biomedical data analysis not merely through their provision of controlled vocabularies but also because they are rich formalized knowledge bases and sources of definitions of domain entities.

## 2 Results

### 2.1 Contribution of axioms in protein-protein interaction prediction

We follow a strategy for the external evaluation of ontologies (Hoehndorf *et al.*, 2012) and apply the method to the task of predicting interactions between proteins and gene–disease associations. Specifically, we intend to test the impact of ontology axioms and ontology meta-data on machine learning applications that rely on ontologies. For this purpose, we use a basic version of GO as the baseline, implement our ontology-based prediction workflows, and evaluate the results. We then compare the performance of ontology-based predictive analysis to the use of GO-Plus in the same workflow and evaluate the results on the same evaluation set. GO-Plus is GO with a large set of formal axioms added that define and constrain GO classes and connect them to classes that are defined in other ontologies (Mungall *et al.*, 2011) (see Section 5.1). Furthermore, we add additional background knowledge in the form of the complete set of axioms in biomedical ontologies that are explicitly used in the GO-Plus axioms, and evaluate their impact on predictive performance.

Since GO-Plus combines all axioms existing in GO with additional axioms that describe relations to other biomedical ontologies, we expect GO-Plus in combination with the axioms and the meta-data of other ontologies to improve predictive performance. We apply GO and GO-Plus to the task of predicting protein-protein interactions (PPIs), and to account for possible differences between taxa in predicting PPIs, we evaluate our hypothesis on human, yeast and Arabidopsis proteins and their interactions (see Section 5.2).

To predict PPIs using GO and GO-Plus, we first assign GO functions to human, yeast and Arabidopsis proteins based on their annotations in the GOA database (Huntley *et al.*, 2015). We then apply the Onto2Vec method (Smaili *et al.*, 2018a), using either GO or GO-Plus as background knowledge, to obtain ontology embeddings of the proteins (see Section 5.2). An ontology embedding is a function that maps entities from an ontology (and its annotations) into an *n*-dimensional vector space (Smaili *et al.*, 2018b), and Onto2Vec encodes for ontology-based annotations of entities together with all the axioms in the ontology (Smaili *et al.*, 2018a). This workflow generates features for proteins based on the same set of GO annotations but utilizes different sets of axioms, and therefore allows us to evaluate the contribution of the ontology axioms to predictions based on these features.

We use the generated features to predict PPIs in two different ways: first, we calculate the cosine similarity between pairs of protein feature vectors (generated through Onto2Vec), and, second, we train a four-layer fully connected neural network on pairs of vectors, and use a sigmoid output to obtain a prediction confidence score (Onto2Vec-NN). We evaluate the results of both prediction methods. Figure 1 shows the ROC curves for PPI prediction for GO and GO-Plus using both Onto2Vec (cosine similarity) and Onto2Vec-NN (neural network) for human, yeast and Arabidopsis Thaliana. Table 1 shows the corresponding AUC values for PPI prediction.

**Table 1.**
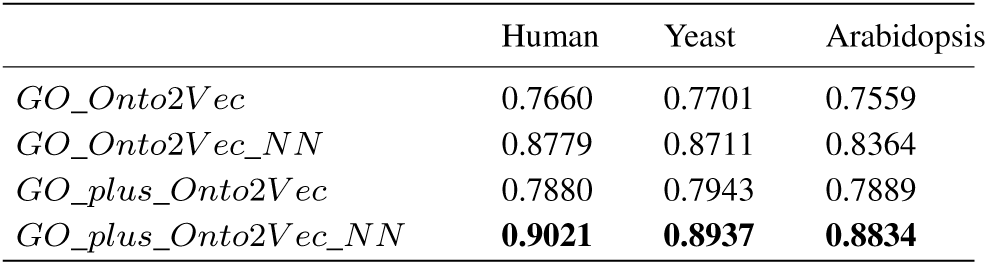
AUC values of ROC curves for PPI prediction for GO-Plus and GO using Onto2Vec (cosine similiarity) and Onto2Vec-NN (neural network).

|  | Human | Yeast | Arabidopsis |
| --- | --- | --- | --- |
| *GO_Onto2Vec* | 0.7660 | 0.7701 | 0.7559 |
| *GO_Onto2Vec-NN* | 0.8779 | 0.8711 | 0.8364 |
| *GO-plus_Onto2Vec* | 0.7880 | 0.7943 | 0.7889 |
| *GO-plus_Onto2Vec-NN* | **0.9021** | **0.8937** | **0.8834** |

**Fig. 1:**
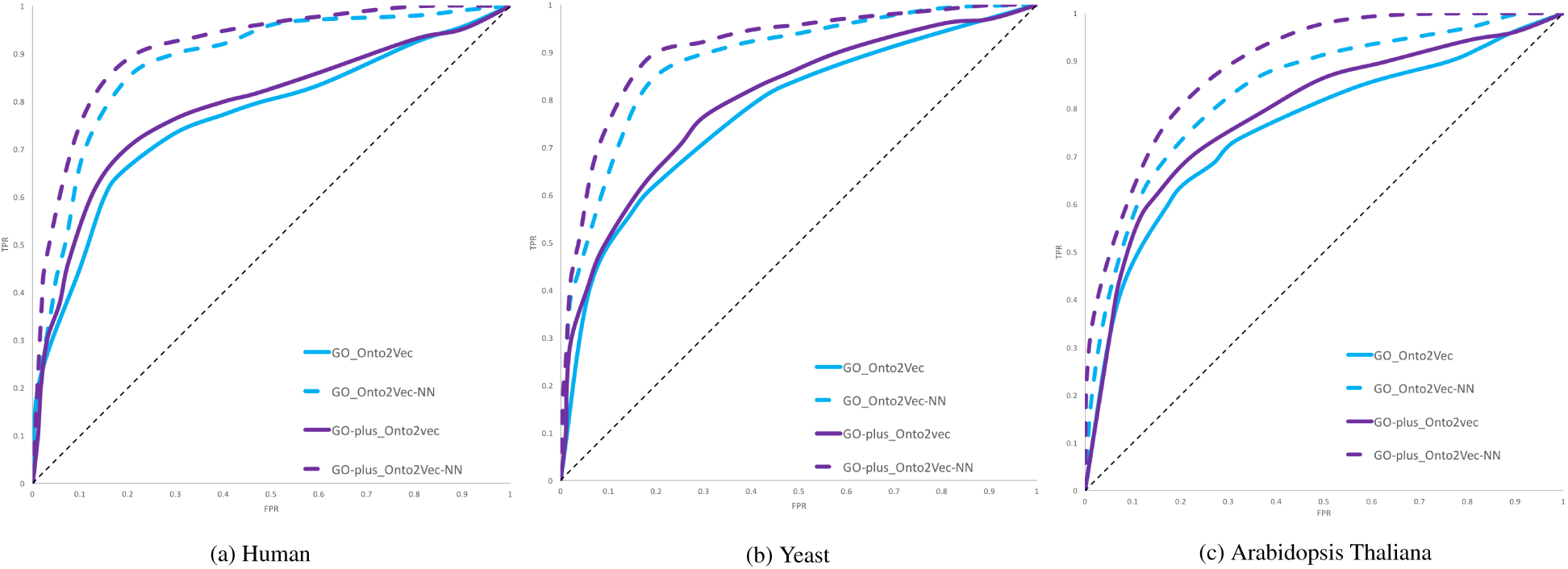
ROC curves for PPI prediction using GO and GO-Plus based on Onto2Vec and Onto2Vec-NN for human, yeast, and Arabidopsis Thaliana.

Our results show that the PPI prediction performance obtained from feature vectors generated using GO-Plus (and the rich set of axioms it contains) outperforms the predictions obtained from using GO axioms alone, both in the unsupervised model (Onto2Vec) and the supervised model (Onto2Vec-NN). The improvement in predictive performance is significant for the Onto2vec prediction based on cosine similarity (*p* = 0.021 for human, *p* = 0.034 for yeast, *p* = 0.027 for Arabidopsis; Mann-Whitney U test), and significant for human and Arabidopsis in the neural network based models (*p* = 0.047 for human, *p* = 0.061 for yeast, *p* = 0.039 for Arabidopsis; Mann-Whitney U test).

GO-Plus uses axioms from many biomedical ontologies but only includes small parts of these ontologies; we hypothesize that the axioms in the ontologies that are referenced in GO-Plus can contribute additional background knowledge that may further improve data analysis. Therefore, we evaluate the individual contribution of each of the ontologies used in GO-Plus axioms, i.e., we individually evaluate the axioms in the Chemical Entities of Biological Interest (ChEBI) ontology (Degtyarenko *et al.*, 2007), the Plant Ontology (PO) (Jaiswal *et al.*, 2005), the Cell type Ontology (CL) (Bard *et al.*, 2005), the Phenotype and Trait Ontology (PATO) (Gkoutos *et al.*, 2005, 2017), the Uberon ontology (Mungall *et al.*, 2012), the Sequence Ontology (SO) (Eilbeck *et al.*, 2005), the Fungal Gross Anatomy Ontology (FAO), the Ontology of Biological Attributes (OBA), the NCBI organismal classification (NCBITaxon), the Common Anatomy Reference Ontology (CARO) (Haendel *et al.*, 2008) and the Protein Ontology (PR) (Natale *et al.*, 2010) (a detailed description of each ontology can be found in Section 5.1).

We repeat the same workflow as before to generate features: representation of GO annotations of the proteins in human, yeast, and Arabidopsis, and representation learning with Onto2Vec using GO-Plus as background knowledge; in each experiment we limit the axioms in GO-Plus to those that contain a reference to a particular ontology. We then again apply Onto2Vec to generate features and predict PPIs through cosine similarity or using a neural network (Onto2Vec-NN) on human, yeast and Arabidopsis.

The AUC values for the PPI prediction using GO-Plus but limited to the axioms that refer a particular ontology are shown in Table 2. We observe that most of the inter-ontology axioms generally improve the predictive performance, with ChEBI contributing the most to improving PPI prediction and PATO improving the least (even decreasing the performance in some cases). The PO is a plant-specific domain ontology and improves predictive performance mainly when predicting PPIs in Arabidopsis, as can be expected.

**Table 2.** AUC values of the ROC curves for PPI prediction showing the contribution of the GO-Plus axioms corresponding to each ontology for human, yeast and Arabidopsis Thaliana. The improvement (blue)/ decrease (red) in performance of each ontology compared to GO is shown between parentheses.

|  | Human |  | Yeast |  | Arabidopsis |  |
| --- | --- | --- | --- | --- | --- | --- |
|  | Onto2Vec | Onto2Vec_NN | Onto2Vec | Onto2Vec_NN | Onto2Vec | Onto2Vec_NN |
| GO (Baseline) | 0.7660 | 0.8779 | 0.7701 | 0.8731 | 0.7559 | 0.8364 |
| ChEBI | 0.7899 (+0.0239) | 0.8914 (+0.0135) | 0.7911 (+0.0210) | 0.8851 (+0.0120) | 0.7703 (+0.0144) | 0.8518 (+0.0154) |
| PO | 0.7752 (+0.0092) | 0.8776 (-0.0003) | 0.7761 (+0.0060) | 0.8741 (+0.0010) | 0.7671 (+0.0112) | 0.8469 (+0.0105) |
| CL | 0.7743 (+0.0083) | 0.8810 (+0.0031) | 0.7819 (+0.0118) | 0.8764 (+0.0033) | 0.7612 (+0.0053) | 0.8371 (+0.0007) |
| PATO | 0.7657 (-0.0003) | 0.8768 (-0.0011) | 0.7707 (+0.0006) | 0.8736 (+0.0005) | 0.7563 (+0.0004) | 0.8380 (+0.0016) |
| UBERON | 0.7780 (+0.0120) | 0.8826 (+0.0047) | 0.7824 (+0.0123) | 0.8781 (+0.0050) | 0.7645 (+0.0086) | 0.8407 (+0.0043) |
| SO | 0.7747 (+0.0087) | 0.8812 (+0.0033) | 0.7763 (+0.0062) | 0.8790 (+0.0059) | 0.7609 (+0.0050) | 0.8375 (+0.0011) |
| FAO | 0.7660 (+0) | 0.8782 (+0.0003) | 0.7712 (+0.0011) | 0.8739 (+0.0008) | 0.7544 (-0.0015) | 0.8368 (+0.0004) |
| OBA | 0.7797 (+0.0137) | 0.8831 (+0.0052) | 0.7874 (+0.0173) | 0.8803 (+0.0071) | 0.7561 (+0.0002) | 0.8371 (+0.0007) |
| CARO | 0.7872 (+0.0212) | 0.8842 (+0.0063) | 0.7881 (+0.0180) | 0.8811 (+0.0080) | 0.7623 (+0.0064) | 0.8503 (+0.0139) |
| PR | 0.7674 (+0.0014) | 0.8784 (+0.0005) | 0.7834 (+0.0130) | 0.8781 (+0.0050) | 0.7669 (+0.0110) | 0.8490 (+0.0126) |
| NCBITaxon | 0.7876 (+0.0216) | 0.8891 (+0.0112) | 0.7892 (+0.0191) | 0.8834 (+0.0103) | 0.7634 (+0.0075) | 0.8479 (+0.0115) |

As a further experiment, we combine all ontologies, i.e., we add the complete set of axioms from each referenced ontology to the axioms of GO-Plus so that the background knowledge in the referenced ontology becomes available to Onto2Vec as well, and then apply our feature learning and prediction workflow. The AUCs for predicting PPIs based on this comprehensive set of ontologies are shown in Table 3. We observe a similar performance to using only the ontology-specific axioms in GO-Plus.

**Table 3.**
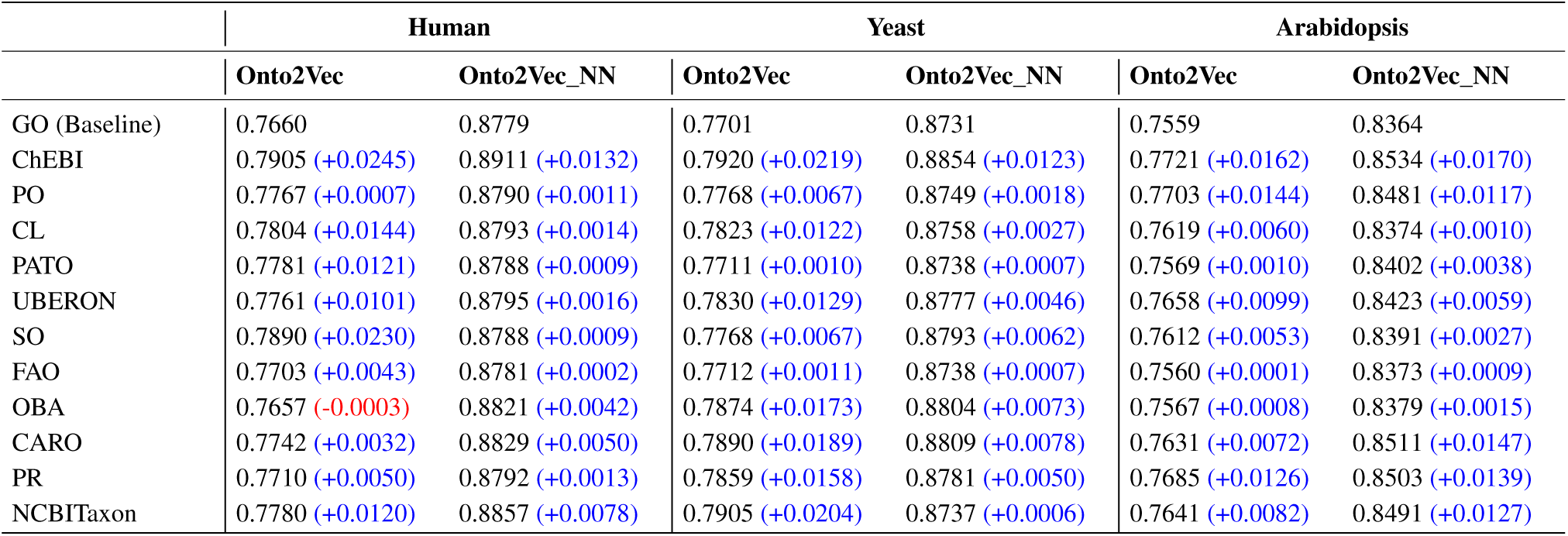
AUC values of the ROC curves for PPI prediction for each external ontology in GO-Plus using Onto2Vec and Onto2Vec-NN. Each prediction method uses all logical axioms from GO, all logical axioms from the referenced ontology, and all GO-Plus axioms describing relations between GO and the given ontology. The improvement (blue)/ decrease (red) in performance of each ontology compared to GO is shown between parentheses.

As a final experiment, we replace Onto2Vec with OPA2Vec to evaluate the contribution of ontology meta-data such as labels, synonyms, and definitions, to their predictive performance (see Section 5.2). We again add each ontology that is referenced in a GO-Plus axiom to the axioms of GO-Plus, this time also including the meta-data (in the form of annotation axioms) of GO-Plus and the referenced ontology. OPA2Vec (pre-trained on the PubMed corpus) can encode both the axioms and meta-data of ontologies and observing the difference to the performance of Onto2Vec can therefore help to evaluate if – and how much – the labels, definitions, and other meta-data contribute.

We again predict PPIs in two different ways: calculating the cosine similarity between the obtained protein feature vectors (referred to as OPA2Vec in the results table) and using the feature vectors to train a neural network for PPI prediction (referred to as OPA2Vec-NN in the results table). Table 4 shows the predictive performance in comparison to using GO. We find that the additional meta-data does, in general, not improve predictive performance; on the contrary, the predictive performance drops markedly when adding the meta-data in several ontologies, most notably PATO and ChEBI.

**Table 4.** AUC values of the ROC curves for PPI prediction for different external ontologies in GO-Plus using OPA2Vec and OPA2Vec-NN. Each prediction method uses the meta-data encoded in GO as well as the meta-data from the external ontologies. In each model, all logical axioms and annotation properties from GO, all logical axioms and all annotation properties from the external ontology, and all GO-Plus inter-ontology axioms are included. The improvement (blue) / decrease (red) in performance of each ontology compared to GO is shown between parentheses.

|  | Human |  | Yeast |  | Arabidopsis |  |
| --- | --- | --- | --- | --- | --- | --- |
|  | OPA2Vec | OPA2Vec_NN | OPA2Vec | OPA2Vec_NN | OPA2Vec | OPA2Vec_NN |
| GO (Baseline) | 0.8727 | 0.9033 | 0.8512 | 0.8891 | 0.8613 | 0.8903 |
| ChEBI | 0.8571 (-0.0156) | 0.8801 (-0.0232) | 0.8411 (-0.0101) | 0.8823 (-0.0068) | 0.8601 (-0.0012) | 0.8880 (-0.0023) |
| PO | 0.8680 (-0.0047) | 0.8824 (-0.0209) | 0.8439 (-0.0073) | 0.8720 (-0.0171) | 0.8632 (+0.0019) | 0.8908 (+0.0005) |
| CL | 0.8811 (+0.0084) | 0.9037 (+0.0004) | 0.8561 (+0.0049) | 0.8891 (+0) | 0.8614 (+0.0001) | 0.8899 (-0.0004) |
| PATO | 0.8562 (-0.0165) | 0.8711 (-0.0322) | 0.8369 (-0.0143) | 0.8696 (-0.0195) | 0.8544 (-0.0069) | 0.8860 (-0.0043) |
| UBERON | 0.8714 (-0.0013) | 0.9033 (+0) | 0.8514 (+0.0002) | 0.8890 (-0.0001) | 0.8615 (+0.0002) | 0.8904 (+0.0001) |
| SO | 0.8711 (-0.0016) | 0.9028 (-0.0005) | 0.8509 (-0.0003) | 0.8879 (-0.0012) | 0.8610 (-0.0003) | 0.8891 (-0.0012) |
| FAO | 0.8709 (-0.0018) | 0.9011 (-0.0022) | 0.8510 (-0.0002) | 0.8882 (-0.0009) | 0.8594 (-0.0019) | 0.8892 (-0.0041) |
| OBA | 0.8774 (+0.0047) | 0.9033 (+0) | 0.8541 (+0.0029) | 0.8897 (+0.0006) | 0.8600 (-0.0013) | 0.8892 (-0.0011) |
| CARO | 0.8808 (+0.0081) | 0.9037 (+0.0004) | 0.8588 (+0.0076) | 0.8900 (+0.0009) | 0.8617 (+0.0004) | 0.8894 (-0.0009) |
| PR | 0.8829 (+0.0102) | 0.9041 (+0.0008) | 0.8590 (+0.0078) | 0.8917 (+0.0026) | 0.8623 (+0.0010) | 0.8911 (+0.0008) |
| NCBITaxon | 0.8704 (-0.0023) | 0.9031 (-0.0002) | 0.8508 (-0.0004) | 0.8886 (-0.0005) | 0.8611 (-0.0002) | 0.8907 (+0.0004) |

### 2.2 Gene–disease association prediction using GO-Plus

In the first part of our analysis we apply GO and GO-Plus to the task of predicting PPIs. Although we utilize PPI datasets from different species for the evaluation in order to generalize our results, it is nevertheless limited to prediction of PPIs and it is unclear if our results also hold for other types of predictive analysis.

We extend our analysis to the evaluation of predicting gene–disease associations based on phenotype similarity (Hoehndorf *et al.*, 2011). While GO is not a phenotype ontology, it is used in the axioms that make up most phenotype ontologies (Gkoutos *et al.*, 2017). We use the cross-species phenotype ontology PhenomeNET (Hoehndorf *et al.*, 2011; Rodríguez-García *et al.*, 2017), which relies on the GO for defining phenotypes, and replace the GO in PhenomeNET with GO-Plus.

We annotate genes with mouse phenotypes from the Mouse Genome Informatics (MGI) (Blake *et al.*, 2017) database as well as disease phenotypes from the Human Phenotype Ontology (HPO) (Köhler *et al.*, 2017) database, and apply Onto2Vec and Onto2Vec-NN (Smaili *et al.*, 2018a) to encode these phenotypes and the axioms in PhenomeNET as feature vectors (more details on the gene–phenotype and disease– phenotype datasets can be found in Section 5.2). We then predict gene–disease associations or mouse models of human disease based on either cosine similarity or a neural network using both Onto2Vec and OPA2Vec. We report the results in Figure 2 and Table 5. The results show that the additional information that GO-Plus provides can significantly improve the overall prediction performance of PhenomeNET in predicting human gene–disease associations and mouse models of human disease ((*p* = 0.0411 and *p* = 0.0254, OPA2Vec, Mann-Whitney U test).

**Table 5.** AUC values of ROC curves for gene–disease prediction using PhenomeNET and when replacing GO in PhenomeNET with GO-Plus.

|  | Human | Mouse |
| --- | --- | --- |
| *Phenomenet + GO_Cos* | 0.7841 | 0.8431 |
| *Phenomenet + GO_NN* | 0.8461 | 0.9141 |
| *Phenomenet + GO - plus_Cos* | 0.7990 | 0.8507 |
| *Phenomenet + GO - plus_NN* | 0.8532 | 0.9182 |
| *Phenomenet + GO - plus + metadata_Cos* | 0.8013 | 0.8672 |
| *Phenomenet + GO - plus + metadata_NN* | **0.8761** | **0.9204** |

**Fig. 2:**
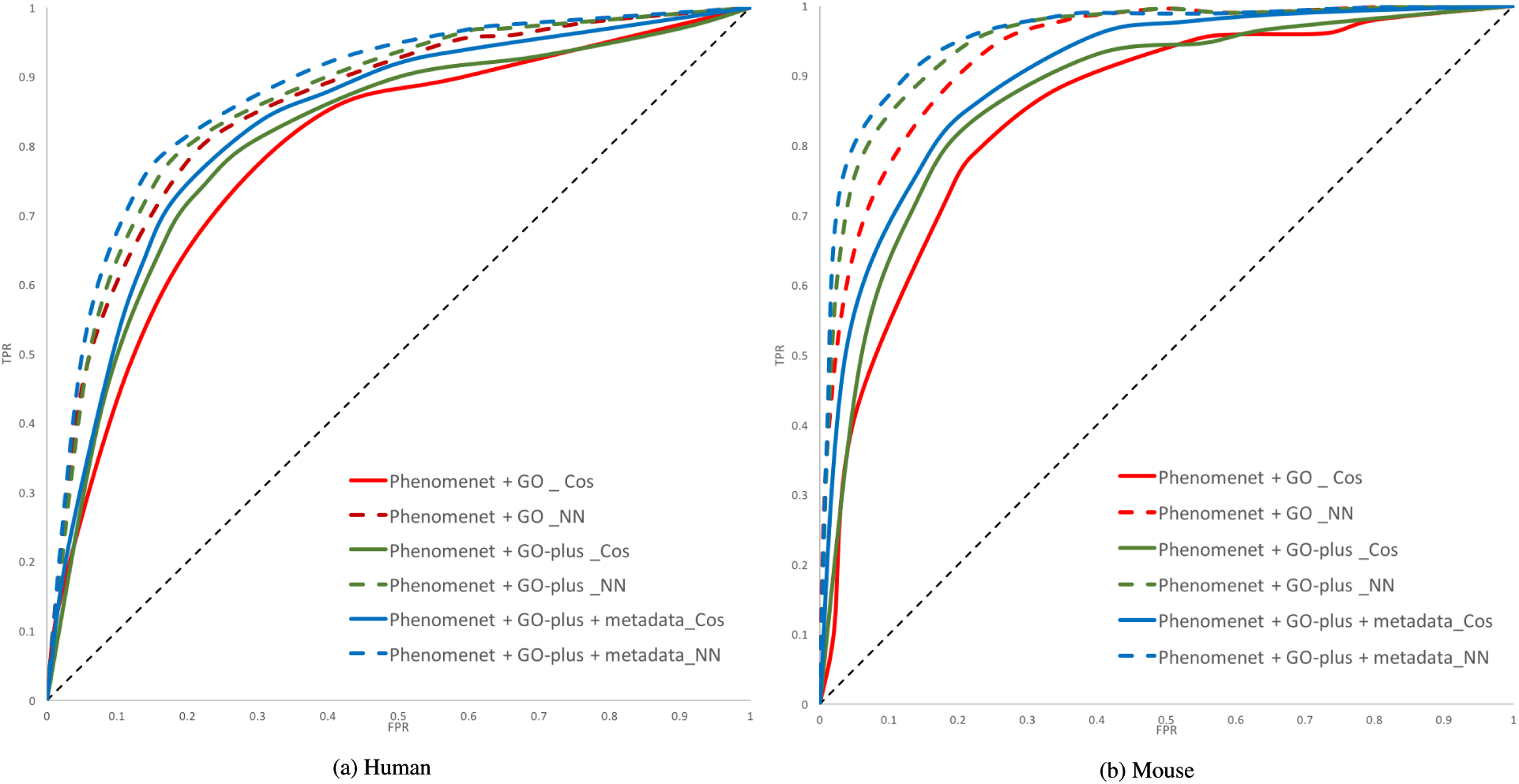
ROC curves for gene–disease prediction comparing PhenomeNET with GO (PhenomeNET + GO) to PhenomeNET with GO-Plus (PhenomeNET + GO-plus) using Onto2Vec and to PhenomeNET with GO-Plus with the metadata included (PhenomeNET + GO-plus + metadata) using OPA2Vec with cosine similarity (Cos) and with a neural network (NN) for human gene–disease associations and mouse models of human disease.

## 3 Discussion

We developed a method to evaluate the contribution of ontology axioms to computational analysis of biomedical data. We use two feature learning methods which are generic and data-driven, and encode for a large set of information contained in ontologies. Our choice is motivated by the desire to avoid potential biases. However, our evaluation is naturally limited to the choice of the two methods (Onto2Vec and OPA2Vec) as well as the application to the prediction of PPIs and gene–disease associations, and the results may change with different application domains.

Nevertheless, our study allows us to draw several conclusions. First, our results demonstrate that including ontology axioms generally adds background knowledge that can significantly improve prediction tasks. Furthermore, our results can be used to improve the axioms as well as textual definitions and labels in existing ontologies. For example, we find that the axioms in ChEBI contribute significantly to the prediction of PPIs because ChEBI axioms reveal relations between GO classes that are associated with the same chemical entities but that are not directly related in the GO hierarchy. Axioms may also add noise to a prediction if they are not well aligned with the prediction task. For example, axioms in the PATO ontology, despite PATO being significantly smaller in size than ChEBI, do not improve or even decrease performance across several applications; furthermore, axioms from the PO only contribute to predicting PPIs in Arabidopsis but not other taxa since PO contains plant-specific domain knowledge.

We also find some evidence that there can be a performance difference when incorporating ontology meta-data into the data analysis. For example, when the OWL annotation axioms of ChEBI are included, the overall PPI prediction performance drops; the labels and definitions in ChEBI often consist of chemical formulas and other properties expressed in symbols or in a mathematical form (e.g., synonyms such as ‘(5Z,8Z,11Z,13E,15R)-15-hydroxyicosa-5,8,11,13-tetraenoic acid’ which are not well represented in literature and therefore not exploited well by our methods. Adding the meta-data (labels, definitions, synonyms, etc.) of the PATO ontology consistently decreases predictive performance across all our applications; a possible explanation for this observation is that the labels and definitions in PATO are not well aligned with any of the tasks we intend to perform; our approach provides a quantitative measure that can be used to improve the PATO definitions and labels for our tasks if this is deemed desirable by the PATO developers.

## 4 Conclusions

We evaluated the contribution of axioms in biomedical ontologies towards predictive analysis methods and found that the background knowledge ontologies provide can significantly improve data analysis and machine learning. Our results have major implications on the further development of knowledge bases and ontologies in the life sciences, in particular as machine learning methods are more frequently applied across the life sciences. Our findings clearly motivate the need for further development, and the systematic, application-driven evaluation and improvement, of formal axioms in biomedical ontologies; and our findings demonstrate this need exists broadly across all areas of biology in which ontologies are applied, not just for a single ontology.

## 5 Materials and Methods

### 5.1 Ontologies

#### GO and GO-Plus

We downloaded the Gene Ontology (GO) (Ashburner *et al.*, 2000) in Web Ontology Language (OWL) (Grau *et al.*, 2008) format from http://www.geneontology.org/ontology/ on April 14, 2018. This version of GO contains 107,762 logical axioms. We also downloaded the GO protein annotations from the UniProt-GOA website (http://www.ebi.ac.uk/GOA) on Dec 2, 2018. All associations with evidence code IEA were filtered, which results in a total of 3,474,539 associations for 749,938 unique proteins.

GO-Plus (downloaded from http://purl.obolibrary.org/obo/go/extensions/go-plus.owl) is an extension of GO that contains, in addition to all the logical axioms of GO, additional inter-ontology axioms that describe relations between GO classes and other external biomedical ontologies, in particular: ChEBI (The Chemical Entities of Biological Interest ontology) (Degtyarenko *et al.*, 2007), PO (The Plant Ontology) (Jaiswal *et al.*, 2005), CL (The Cell Ontology) (Bard *et al.*, 2005), PATO (Phenotype and Trait Ontology) (Gkoutos *et al.*, 2005, 2017), the Uberon ontology (Mungall *et al.*, 2012), SO (The Sequence Ontology) (Eilbeck *et al.*, 2005), FAO (Fungal gross anatomy), OBA (Ontology of Biological Attributes), NCBITaxon (NCBI organismal classification), CARO (Common Anatomy Reference Ontology) (Haendel *et al.*, 2008) and PR (Protein Ontology) (Natale *et al.*, 2010). Table 6 summarizes the number of axioms in GO-Plus describing relations to each of these ontologies and shows an example of such axioms for each ontology.

**Table 6.**
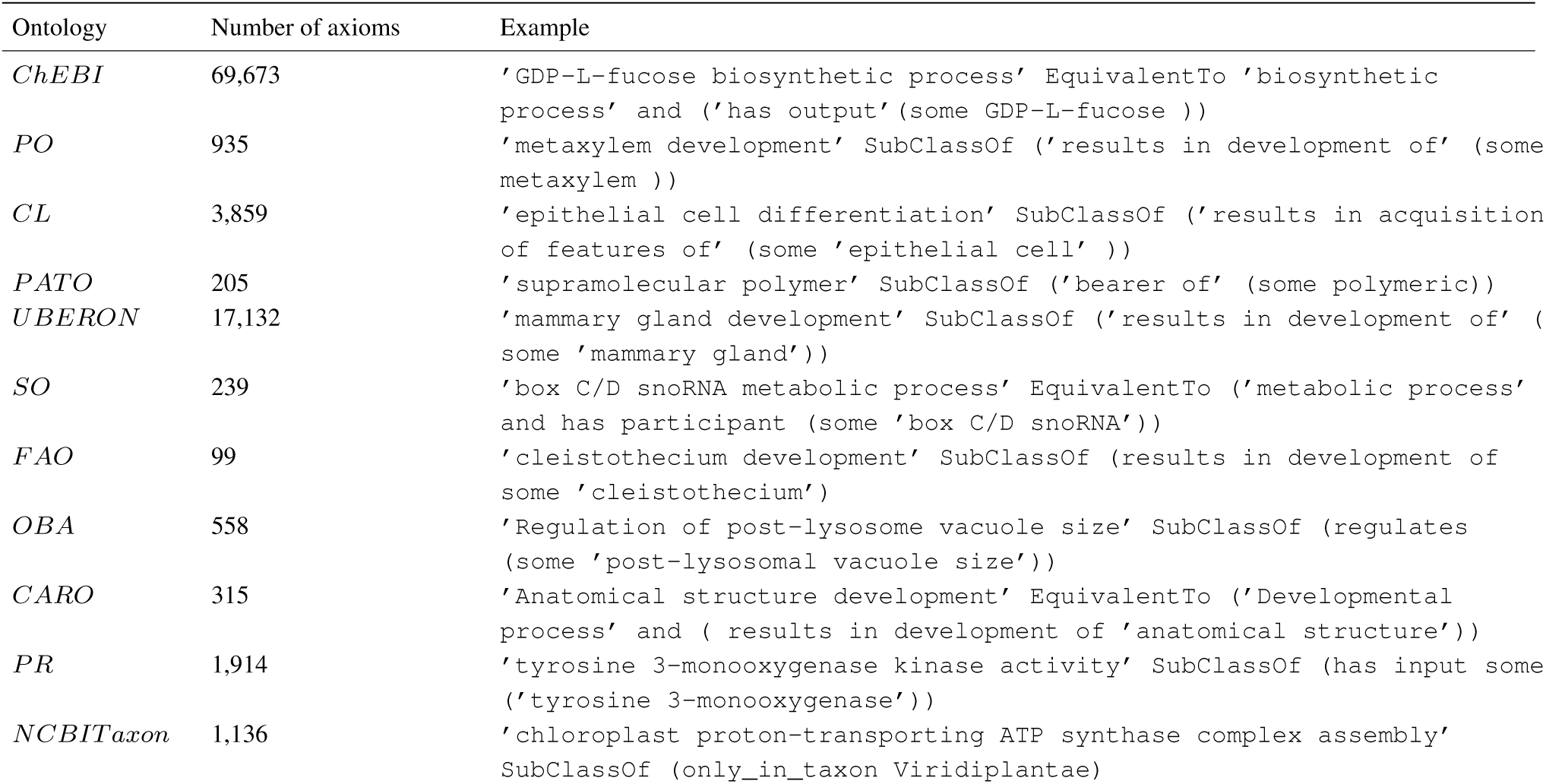
Number of inter-ontology axioms (with an example) in GO-Plus corresponding to each external ontology.

#### The ChEBI Ontology

We downloaded ChEBI in the OWL format from http://purl.obolibrary.org/obo/chebi.owl on April 26, 2018. The ChEBI ontology formally describes relations between molecular entities, in particular small chemical compounds (Degtyarenko *et al.*, 2007). It contains a total of 432,822 logical axioms and 92,015 classes.

#### The Plant Ontology (PO)

We downloaded the OWL version of PO from http://purl.obolibrary.org/obo/po.owl on April 26, 2018. This version of PO contains 4,835 axioms and 1,649 classes. PO provides a formal description of the vocabulary related to external and internal plant anatomy and plant development phases. It is mainly used to associate plant structures and development to gene expression and phenotype data (Cooper *et al.*, 2013).

#### The Cell Type Ontology (CL)

We downloaded CL in OWL from http://purl.obolibrary.org/obo/cl.owl on April 26, 2018. CL contains 17,958 axioms and 3,862 classes. It is an ontology that describes cell types for major animal and plant organisms (Bard *et al.*, 2005).

#### Phenotype and Trait Ontology (PATO)

The OWL version of PATO was downloaded from April 26, 2018 from http://purl.obolibrary.org/obo/pato.owl. This version contains 5,644 logical axioms and 2,251 different classes. PATO provides a systematic description of phenotypes through the concepts and relationships defined by its axioms (Gkoutos *et al.*, 2005).

#### Uberon Ontology

We downloaded the Uberon ontology on April 26, 2018 from http://purl.obolibrary.org/obo/uberon.owl. This OWL version of Uberon contains 65,067 logical axioms and 9,866 classes. Uberon is a multi-species anatomy ontology that describes anatomical structures across multiple species through manually-curated cross-references (Mungall *et al.*, 2012).

#### Sequence Ontology (SO)

We obtained the SO ontology from http://purl.obolibrary.org/obo/so.owl on November 25, 2018. This version of SO contains 5,443 logical axioms and 2,2234 classes. The SO consists of a set of classes and relations that describe the parts of a genomic annotation (Eilbeck *et al.*, 2005).

#### Fungal Gross Anatomy Ontology (FAO)

We downloaded the FAO ontology on November 25, 2018 from http://purl.obolibrary.org/obo/fao.owl. The OWL version of FAO contains 155 axioms and 105 classes. The FAO describes the anatomy of fungi through a set of controlled vocabulary.

#### Ontology of Biological Attributes (OBA)

We downloaded the OBA ontology on November 25, 2018 from http://purl.obolibrary.org/obo/oba.owl. This ontology contains 73,377 axioms and 27,365 classes. OBA provides a collections of biological attributes.

#### NCBI organismal classification (NCBITaxon)

We obtained the NCBITaxon ontology from http://purl.obolibrary.org/obo/ncbitaxon.owl. This OWL version contains 3,653,676 axioms and 1,826,669 classes. This ontology provides a formal classification of different organisms (Federhen, 2011).

#### Commom Anatomy Reference Ontology (CARO)

The CARO ontology was obtained on http://purl.obolibrary.org/obo/caro.owl on November 25, 2018. This version contains 209 axioms and 158 classes. The CARO serves as a template to unify the structure of anatomy ontologies (Haendel *et al.*, 2008).

#### Protein Ontology (PR)

We downloaded the PR ontology from http://purl.obolibrary.org/obo/pro_reasoned.owl on November 4, 2018. This ontology contains 1,312,362 axioms and 400,923 classes. The PR ontology formally represents protein-related entities and their relations at different levels of specificity(Natale *et al.*, 2010).

#### PhenomeNet Ontology

We downloaded the PhenomeNET ontology (Hoehndorf *et al.*, 2011; Rodríguez-García *et al.*, 2017) in OWL format from the AberOWL repository http://aber-owl.net (Hoehndorf *et al.*, 2015a) on February 21, 2018. PhenomeNET is a cross-species phenotype ontology that combines phenotype ontologies, anatomy ontologies, GO, and several other ontologies in a formal manner (Hoehndorf *et al.*, 2011).

### 5.2 Evaluation Datasets

#### Protein-protein interactions (PPI)

To evaluate our work, we predict PPI on three different organisms: human, yeast, and *Arabidopsis thaliana*. The datasets for all three organisms were obtained from the STRING database (Szklarczyk *et al.*, 2017)(http://string-db.org).The human dataset contains 19,577 proteins and 11,353,057 interactions, the yeast dataset contains 6,392 proteins and 2,007,135 interactions, while the Arabidopsis dataset contains 10,282,070 interactions for 13,261 proteins.

#### Gene–disease associations

To further evaluate our method, we predict gene–disease associations. The first dataset used in this experiment is the mouse phenotype annotations obtained from the Mouse Genome Informatics (MGI) database (Smith and Eppig, 2015) on February 21, 2018 with a total of 302,013 unique mouse phenotype annotations. The second dataset used for this experiment is the disease to human phenotype annotations obtained on February 21, 2018 from the Human Phenotype Ontology (HPO) database (Robinson *et al.*, 2008). We limited our analysis to the OMIM diseases only which resulted in a total of 78,208 unique disease-phenotype associations. To validate our prediction, we used the `MGI_DO.rpt` file from the MGI database to obtain 9,506 mouse gene-OMIM disease associations and 13,854 human gene-OMIM disease associations. To map mouse genes to human genes we used the `HMD_HumanPhenotype.rpt` file from the MGI database.

### Analysis algorithms

Our analysis is based on prediction results obtained using embeddings of biological entities (proteins, genes, diseases) obtained from ontologies using two tools: Onto2Vec (Smaili *et al.*, 2018a) and OPA2Vec (Smaili *et al.*, 2018b). The obtained embeddings are then trained using a neural network to make predictions.

#### Onto2Vec

Onto2Vec (Smaili *et al.*, 2018a) is a method that uses ontologies to obtain embeddings of ontology classes and the entities they annotate. Onto2Vec uses two main information sources: First, it used all logical axioms describing the structure of an ontology including both asserted axioms of an ontology as well as inferred axioms using a semantic reasoner. Second, it uses the known ontology-based associations of biological entities (e.g. protein-GO associations). These two pieces of information form a corpus of text used to train word2vec (Mikolov *et al.*, 2013b,a) and obtain the embeddings.

#### OPA2Vec

OPA2Vec (Smaili *et al.*, 2018b) is also a tool used to obtain embeddings of biological entities from ontologies. In addition to using logical axioms, OPA2Vec also uses annotation property axioms from the ontology meta-data. These annotation axioms use natural language to describe different properties of the ontology classes (labels, descriptions, synonyms, etc.) and they, therefore, form a rich corpus of text for word2vec. To provide the word2vec model with some background knowledge on the ontology concepts described by the annotation properties, OPA2Vec pre-trains the model on a corpus of biomedical text (PubMed by default). Entity-class annotations are also used an additional source of information to produce the ontology-based embeddings of biological entities.

#### Cosine similarity

One way to perform prediction tasks using ontology-based embeddings is by calculating the similarity between each pair of vectors and using the obtained similarity as a confidence score to predict whether two entities are associated or not. To do so, we use cosine similarity as a similarity measure between the obtained vectors. The cosine similarity *cos_sim_* between two vectors *A* and *B* is calculated as
f

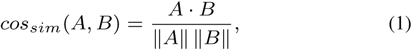

where *A • B* is the dot product of *A* and *B*.

#### Neural Network

To optimize our prediction models (PPI and gene–disease associations predictions), we train a neural network using the obtained embeddings from both Onto2Vec and OPA2Vec. Limited grid search has been performed to select a suitable neural network for our predictions based on suggested guidelines (Hunter *et al.*, 2012). The chosen neural network is a feed-forward network with two hidden layers of 800 and 200 neurons respectively. The neural network is optimized using binary cross entropy as the loss function.

### Evaluation metrics

We used the ROC (receiver operating characteristic) curve (Yin and Vogel, 2017) along with the AUC (area under ROC curve) as a quantitative measure to assess the performance of each predictive method. For both PPI prediction and gene–disease prediction, the true positive pairs are considered to be the ones available from the STRING network and the `MGI_DO.rpt` file from the MGI database respectively. The negative pairs on the other hand are down-sampled from the set of all unknown associations to form a set of negatives equal in size to the set of positive pairs for both PPI prediction and gene–disease association prediction.

